# A functionally selected *Acinetobacter* sp. phosphoethanolamine transferase gene from the goose fecal microbiome confers colistin resistance in *E. coli*

**DOI:** 10.64898/2025.12.09.693354

**Authors:** Elizabeth Bernate, Yijun Shi, Ezabelle Franck, Terence S. Crofts

**Affiliations:** College of Veterinary Medicine, University of Florida, Gainesville, FL; Department of Animal Sciences, University of Illinois at Urbana-Champaign, Urbana, IL; Department of Biomedical Sciences, Florida State University, Tallahassee, FL; Carl R. Woese Institute for Genomic Biology, University of Illinois at Urbana-Champaign, Urbana, IL; Division of Nutritional Sciences, University of Illinois at Urbana-Champaign, Urbana, IL

## Abstract

Polymyxins are last-resort antibiotics for infections caused by multidrug resistant Gram-negative bacteria such as Enterobacteriaceae, *Pseudomonas aeruginosa* and *Acinetobacter baumannii*. This makes the rise of bacteria exhibiting polymyxin E (colistin) resistance, largely through modification of lipid A moieties, concerning and suggests that it is important to document potential sources of the corresponding resistance genes. This study searched for potential emerging colistin-resistance genes from the environment by investigating a previously performed functional metagenomic selection for colistin resistance of a goose fecal microbiome. We found that the selection captured *Acinetobacter* sp. DNA fragments which all contained *eptA* genes. We confirmed their ability to confer significant colistin resistance in *E. coli via* modification of lipid A in the outer membrane. Furthermore, we found evidence for mobilization of closely related *eptA* genes in *Acinetobacter* strains, marking them as potential *mcr* genes or their precursors. This study highlights the goose fecal microbiome as a potential source for colistin resistance in the environment.

## INTRODUCTION

Polymyxin antibiotics were discovered and isolated in 1947 by Y. Koyama from the soil bacterium *Paenibacillus polymyxa* subspecies *colistinus* (1). As antibiotics, polymyxins were initially disregarded due to their harmful side effects compared to other antibiotic classes but they were eventually adopted into the clinic due to their ability to treat otherwise antibiotic-resistant Gram-negative pathogens. The only commercially available compounds from this class are polymyxin B and polymyxin E (aka colistin) (2). Polymixin B and colistin are both peptide antibiotics and differ by only one amino acid from each other and have similar microbiological activity (3). Polymyxin B is used topically while colistin can be used to treat systemic infections as well (4). Colistin has two different commercially available forms: colistin sulfate as an oral and topical medication and colistimethate which can be administered *via* injection (5). Colistin was first approved and used in the 1950s against otherwise resistant Gram-negative bacterial infections and was praised for its comparatively low level of antibiotic resistance in both a human and veterinary setting (1). In the 1970s, it became less frequently used in human medicine due its side effects of nephrotoxicity and neurotoxicity (6) but was reintroduced into the clinic in the 1990s (7) for treating Cystic Fibrosis-associated Gram-negative infections and colistin now serves as a last line of defense against MDR infections. In contrast to human medicine, colistin has been continuously and extensively used in veterinary medicine (8) for the treatment of diseases by bacteria such as *E. coli* and *Salmonella* in rabbits, poultry, livestock (9) and *Aeromonas* and *Shewanella* in aquaculture (10, 11). In agricultural settings (mainly swine), colistin is or has been given orally through drinking water or feed for multiuse infection prevention and growth promotion (12). Colistin is active against many Gram-negative bacteria and there are concerns that its prophylactic use in agriculture, leading to release of colistin into other environments, contributes to increased selection for bacterial resistance (8, 13).

Colistin is an acylated cyclic polypeptide that is composed of over 30 polymyxins, mainly polymyxin E1 and polymyxin E2, and operates by interacting as a net positively charged structure, binding to the negative phosphates of the lipid A lipopolysaccharide (LPS) of Gram-negative bacteria. Binding of colistin to LPS destabilizes the calcium and magnesium bridges on the outer membrane *via* hydrophobic and electrostatic interaction (14). This destabilization causes release of the LPS and permeabilizes the outer membrane, allowing the antibiotic to pass to the cytosolic membrane through the lipophilic acid-fat chain where it causes disruption of the phospholipid bilayer, resulting in osmotic disparity between the inside and outside of the cell and, eventually, lysis (1, 15). Other mechanisms may be involved in its antibacterial activity, such as inhibition of respiratory enzymes (16) and induction of oxidative stress caused by an imbalance of oxygen reactive species (17).

The dominant mechanism of colistin resistance acts by preventing binding of the drug to its lipid A target. This can occur through the transfer of cationic moieties onto lipid A, either by a phosphoethanolamine transferase (*e.g.,* MCR and EptA) or an aminoarabinose transferase (*e.g.,* ArnT). This outer membrane modification results in an increased density of positive charges on the cell membrane and decreases the binding affinity of colistin (**Figure 1**). Additionally, chromosomal mutations effecting synthesis of lipid A or increased lipid A 4’-phosphatase activity can also cause a decrease in negatively charged phosphates in the outer membrane, ultimately decreasing the electrostatic interaction and affinity of colistin to bind and affect the bacteria (7, 18–20). Additional mechanisms of colistin resistance exist that do not rely on lipid A modification. These include active efflux of colistin and biofilm formation, both of which decrease the amount of colistin that reaches the cytosolic membrane (2, 21, 22).

**Figure 1.**
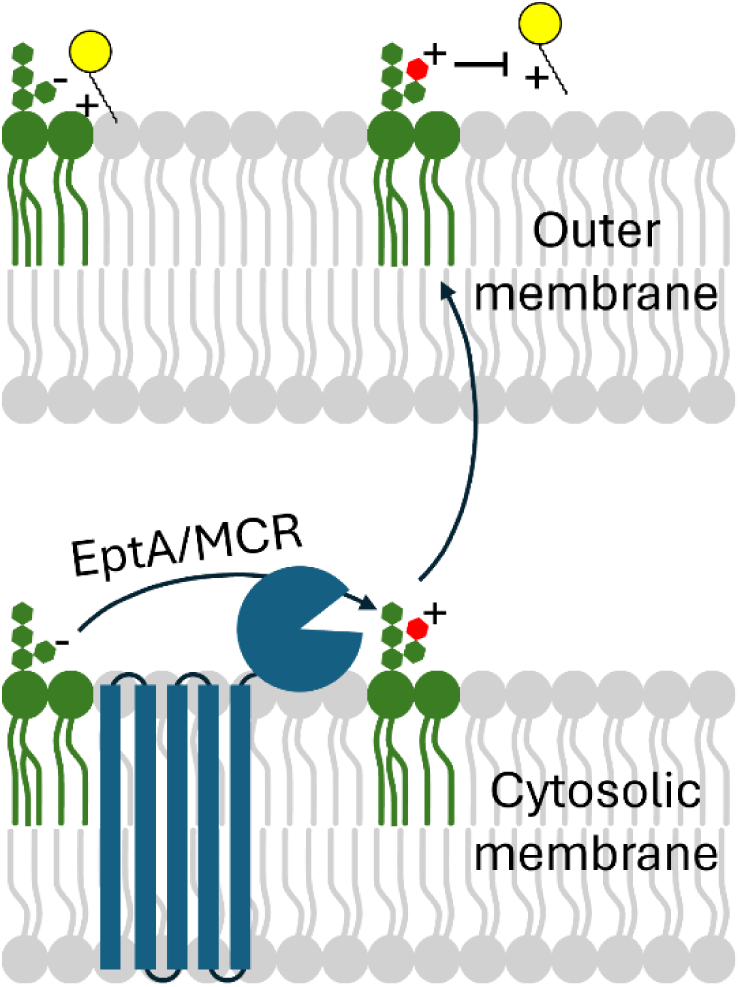
Phosphoethanolamine transferases confer colistin resistance. Membrane-bound enzymes EptA and MCR (shown in blue) transfer a cationic phosphoethanolime group (red) to lipid A (green), changing its electrostatic characteristics. This decreases the ability of colistin, or other polymyxin antimicrobials (yellow), to interact with lipid A, conferring decreased susceptibility.

Colistin resistance can also arise from a regulatory two-component system PmrAB (coupled to PhoP/Q by PmrD) that upregulates lipid A modification operons (6, 23, 24). Acquired resistance can be due to mutations or changes in genomic context of this system that trigger activation of lipid A modification genes. More concerning are mobilized colistin resistance mechanisms. In 2015, as part of an antimicrobial resistance monitoring project in China, a colistin resistant *E. coli* strain was isolated from a pig (strain SHP45) (3). Resistance was determined to be conferred by a gene carried on a plasmid. Because the gene could potentially be mobilized to other bacteria *via* the plasmid it was termed *mcr*-1 for <u>m</u>obilizable <u>c</u>olistin resistance. The *mcr* gene family encode phosphoethanolamine transferases, similar to those encoded by the *eptA* gene family, which confer resistance through charge alteration of lipid A (25–27) (**Figure 1**). Studies have since found evidence for *mcr* genes in *E. coli* isolates from as far back as the 1970s (28) and in a variety of plasmid contexts and bacterial hosts (29) suggesting their spread across environments.

The early emergence of *mcr* and its characteristic of transmissibility suggest that there are homologs in the natural environment yet to be found. This is born-out by the rapid discovery of *mcr* genes 2 through 10 (30). Here, we used a functional metagenomics approach to search for potential colistin-resistance genes from the fecal microbiome of a wild goose. Functional metagenomics is a technique that allows for sequence-naïve, high-throughput identification of genes in an entire microbiome’s metagenome based on their ability to confer antibiotic resistance. Functional metagenomic libraries are prepared by extraction of metagenomic DNA from a microbiome (in our case, a goose fecal pellet), fragmentation of metagenomic DNA, cloning of the random metagenomic DNA fragments into a plasmid, and transformation of an *E. coli* expression host with the plasmid library (**Figure 2**). This *E. coli* library is then spread onto agar plates containing inhibitory concentrations of antibiotics (*e.g.,* 4 µg/ml colistin), and plasmids collected from any colonies that grow under these conditions are sent for sequencing (**Figure 2**). This technique can identify both known and novel antibiotic resistance genes as well as potentially identify evidence of horizontal-transfer of antibiotic-resistance genes (31, 32).

**Figure 2.**
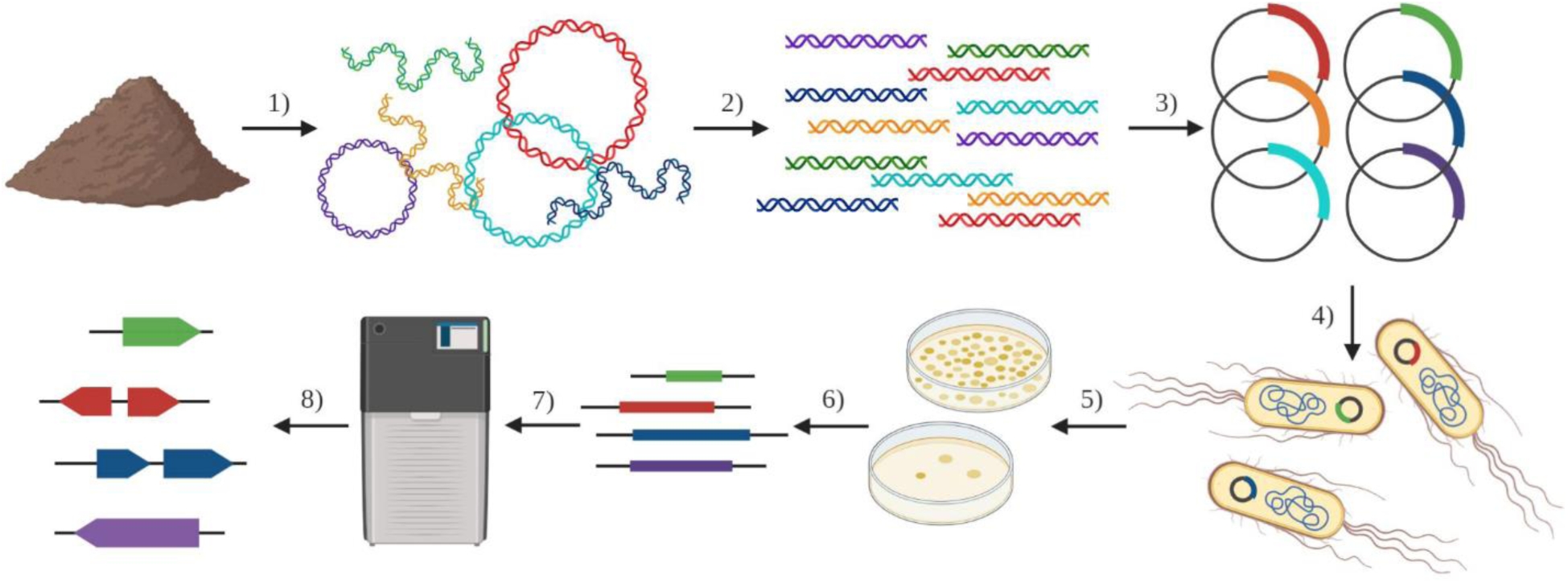
Functional metagenomic library construction and selection. 1) Metagenomic DNA is extracted from a microbiome of interest (such as a goose fecal pellet) and 2) fragmented into ∼2 kb pieces for 3) insertion into plasmid vectors. An *E. coli* host strain is 4) transformed by the plasmid library where potential metagenomic genes are transcribed and translated into proteins. The functional metagenomic library is 5) selected by plating on colistin-containing agar plates. Surviving resistant colonies are collected and 6) resistance-conferring metagenomic DNA fragments are 7) sequenced and 8) potential colistin resistance genes are identified and annotated. (Figure from Crofts *et al.*, 2021).

## METHODS AND MATERIALS

### Bacterial strains, reagents, and chemicals

Cultivation of *E. coli* (DH10B genotype) was generally performed at 35°C, with aeration by shaking at 250 rpm for liquid cultures, unless otherwise specified. Routine cultures were grown in lysogeny broth (Miller) (LB) with 50 μg/ml kanamycin (LB+KAN50) for maintenance of plasmids. Antimicrobial susceptibility testing was carried out using Cation-Adjusted Mueller-Hinton media (MH) (Teknova, 101320-364) with 50 μg/ml kanamycin (MH+KAN50) when appropriate. Plasmids used in this study were derivatives from pZE21 unless otherwise noted(33–35). Bacterial clones were stored in a -80°C freezer as 15% glycerol stocks in LB.

The *mcr*-expressing positive control *E. coli* strain used during antimicrobial susceptibility testing and lipid A mass spectrometry analysis was from the Minimal Antibiotic Resistance Platform (ARP) and a gift from Gerard Wright (Addgene kit #1000000143) (36).

Reagents and chemicals were of molecular biology grade or higher purity. Kanamycin sulfate (VWR Life Science, 75856-68), and colistin sulfate (Sigma Aldrich, C44611G) powders were stored at 4°C in a desiccator while solutions were stored at -20°C as sterile filtered 50 mg/ml stock solutions. Minimal inhibitory concentration test strips for polymyxin B (cat# 920041) and colistin (cat# 921411) were purchased from Liofilchem.

### Functional metagenomic selection

A goose fecal microbiome functional metagenomic library was previously constructed and selected for colistin resistance (31). Briefly, the library was prepared from metagenomic DNA extracted from a Canada goose fecal pellet and encoded ∼27 Gb of metagenomic DNA with an average insert size of ∼2.4 kb. The library was plated onto Mueller-Hinton cation adjusted II (MH) agar plates containing either 4 μg/ml or 8 µg/ml colistin sulfate and incubated overnight at 37°C. Initially, no resistant colonies were observed on the 8 μg/ml agar plate but two were found after an additional incubation at room temperature which were saved for analysis. The agar plates containing 4 µg/mL colistin resulted in ∼360 resistant colonies after incubation overnight. The resistant colonies were collected from the agar plate by resuspending them with a cell spreader in 1 ml of LB (4 μg/ml selection) or picking into LB+KAN50 media and culturing (8 μg/ml colonies). Plasmids were isolated from the resulting slurry or cultures *via* miniprep (New England Biolabs, T1010L) according to manufacturer protocols and stored at - 20°C until sequencing.

### Metagenomic fragment sequencing and processing

Colistin resistance-conferring metagenomic fragments from the pooled slurry of the 4 μg/ml colistin plate colonies were previously sequenced (33). Briefly, the metagenomic inserts were amplified by PCR from the slurry miniprep using Q5 polymerase (New England Biolabs, M0492S) in a 100 μl reaction with 5 ng of DNA as template using primers that amplify from 250 bp either side of the vector mosaic end sequence sites. The thermocycler settings included an initial 30 second hold at 98°C followed by ten cycles of 98°C for 10 seconds and 72°C for 4 minutes. The amplified metagenomic inserts were purified by silica column kit (New England Biolabs, T1030S) and quantified by fluorescence (Promega QuantiFluor, E4871). The resulting purified colistin resistance-conferring amplicons were submitted for sequencing on the PacBio Sequel II platform at the Roy J. Carver Biotechnology Center at the University of Illinois at Urbana-Champaign.

For analysis, the PacBio reads were first processed in Galaxy to remove vector sequences from the reads (37). The trimmed reads were dereplicated using cd-hit-est with a 99% nucleotide identity cut-off and otherwise default settings (38–40). The resulting clusters were sorted by number of reads in the cluster, and the representative read from each was extracted for analysis. Metagenomic fragments from the 16 highest rank clusters were selected for cloning.

### Cloning of metagenomic fragments

The selected metagenomic DNA fragments from the 4 µg/ml functional metagenomic selection were amplified and cloned into a pZE21 derivative (34) with a modified cloning site 8 bp downstream of the vector ribosome binding site (pTSC174) (41). Amplification was carried out using Q5 polymerase with a 2-step PCR protocol (New England Biolabs, M0492S) and inserts were cloned into pTSC174 using NEBuilder HiFi assembly mix (New England Biolabs, E2621L) according to manufacturer protocols. Recombinant plasmid was transformed into chemically competent *E. coli* DH10B cells using heat shock (42), and plasmids from the transformed *E. coli* were verified to have the correct sequences by full plasmid sequencing through Plasmidsaurus. Plasmids extracted from the two *E. coli* clones that grew on the 8 μg/ml colistin plates were sequenced without re-cloning.

### Bioinformatic Analysis

Analysis of the gene contents of the metagenomic inserts was performed by calling open reading frames and annotating the resulting predicted proteins using the Bakta server (43, 44), MetaGeneMark (45), and by using blastp to align the predicted proteins against non-redundant protein sequences (conducted October, 2025) (46, 47). Relative coordinates for the predicted genes were used to construct gene schematics.

The EptA predicted to be encoded on the 8 μg/ml selected metagenomic DNA fragment was compared to representative EptA, EptB, EptC, and MCR protein sequences. UnitProt 50% Reference Clusters (48, 49) were downloaded for EptA, EptB, and EptC. MCR sequences (ARO:3004268) were downloaded from the Comprehensive Antibiotic Resistance Database (CARD) (50, 51). Protein sequences were aligned using the Clustal Omega webserver (52) and the alignment was used to construct a maximum likelihood tree with IQ-Tree (VT+R6 model, 100 replicates) (53, 54). Redundant enzymes and those lacking genus level annotation were trimmed and the tree was visualized with the Interactive Tree of Life (iTOL) program (55).

### Antimicrobial Susceptibility Tests

Agar dilution antimicrobial susceptibility assays were carried out as published in Wiegan *et al.* (56). Briefly, triplicate single colonies from each strain were resuspended in 200 μl of MH+KAN50 broth to reach an OD_600_ at 1 cm of between 0.08 and 1 absorbance units (equivalent to a 0.5 McFarland standard) to produce a bacterial suspension of ∼1×10^8^ colony forming units (cfus) per ml. Triplicate suspensions were diluted 10-fold with MH broth in a 96-well plate. A 48-pin microplate replicator (Fisher Scientific, 05-450-10), sterilized by dipping into 70% ethanol and flame dried, was placed in the wells to pick up 1 µL of each culture and stamped onto agar plates composed of MH+KAN50 and the concentration of colistin being tested (0 µg/ml and 0.03125 μg/ml to 32 µg/ml by 2-fold increments). The agar plates were incubated at 35°C overnight and photographed and analyzed following 20 to 24 hr of growth.

50% inhibitory concentration values (IC_50_) for colistin were determined using broth microdilution assays (56, 57). A 96-well plate was prepared with each well containing 50 μl of MH+KAN50 and colistin. Colistin concentrations varied by two-fold increments across 10 columns, with the highest colistin concentration being 128 μg/ml. Two control volumes did not contain colistin (column 11 as a growth control and column 12 as a sterility control). A 0.5 McFarland standard for each of the tested strains was prepared as before and diluted 100-fold before 50 μl aliquots were added to the antibiotic-containing 96-plate to inoculate the wells. The plate was sealed with a Breathe-Easy membrane (MilliporeSigma, Z380059) and incubated at 35°C with shaking at 250 rpm overnight for 20 hr to 24 hr. Growth was quantified by measuring OD_600_ absorbance on a plate reader (Agilent Biotek, Synergy HTX) and dose-response curves were generated to compute IC_50_ values using Graphpad Prism (version 10.6.1).

Liofilchem colistin and polymyxin test strips were used to find MIC values according to manufacturer instructions. Briefly, colonies of the strains being analyzed were inoculated in MH broth to produce a 0.5 McFarland standard as above. A sterile cotton-tipped swab was dipped into the culture and used to inoculate the full surface of MH+KAN50 agar plates. Antibiotic test strips, one for colistin and one for polymyxin B, were placed on the plates. The cultures were incubated at 35°C for 20 hr to 24 hr, after which they were photographed and analyzed to determine MIC values.

### Mass spectrometry analysis of lipids

Lipid A molecules were extracted from *E. coli* cultures grown overnight at 35°C with shaking at 250 rpm in LB+KAN50 following the sodium acetate method described by Liang *et al.* (58). Cultures (1 ml) were pelleted by centrifugation at 8,000 rcf for 1 minute and supernatants removed. The pellets were resuspended in 400 μl of 100 mM, pH 4 sodium acetate buffer and held at 99°C for 30 minutes with mixing by briefly vortexing every 10 minutes. The samples were brought to room temperature by placing on ice then centrifuged at 8,000 rcf for 5 minutes. The insoluble pellet was washed once with 95% ethanol and lipids were extracted from the washed pellet by addition of 100 μl of a chloroform/methanol/water mix (12:6:1 by volume). Insoluble material in the lipid extract was removed by centrifugation at 5,000 rcf for 5 minutes. The clarified lipid solution was characterized by matrix-assisted laser desorption/ionization time-of-flight mass spectrometry using a Bruker UltrafleXtreme MALDI TOFTOF instrument in negative reflector mode at the University of Illinois at Urbana-Champaign School of Chemical Sciences Mass Spectrometry Laboratory.

## RESULTS

### Characterization of colistin resistance DNA fragments from a functional metagenomic selection of the goose fecal microbiome

We set out to characterize colistin resistance genes from the goose fecal microbiome using functional metagenomic selections. We previously selected a 27 Gb functional metagenomic library prepared from goose fecal metagenomic DNA on agar plates containing 4 μg/ml colistin sulfate and collected approximately 360 resistant colonies from the selection (33). We sequenced the resistance-conferring metagenomic inserts and clustered them at 99% nucleotide identity, resulting in 5576 clusters, the majority of which contained a single read (**Supplemental figure 1**). After stack ranking the clusters by number of reads, we focused on the top 16 largest clusters for analysis which accounted for 68% of all clustered reads (CST1-CST16). We added an additional metagenomic DNA fragment to the analysis that originated from an extended selection of the same library on an agar plate containing 8 μg/ml colistin sulfate (CST17).

We re-cloned a total of 15 metagenomic inserts out of the 16 identified, with clones corresponding to CST1 and CST15 failing the cloning process. We re-sequenced the inserts by long read sequencing and analyzed them by blastn and alignment and found all of them to be closely related to *Acinetobacter* sp. ASP199 (**Supplemental table 1**). The inserts showed a high level of identity to each other (97.62% average nucleotide identity across inserts, **Supplemental figure 2A**) and a high level of clustering (**Supplemental figure 2B**) (with CST12 appearing to have acquired a 3’ truncation). After predicting genes on the metagenomic inserts and annotating the resulting open reading frames it became clear that they all contained a similar genomic construction: one to two predicted regulatory genes, a phosphoethanolamine transferase gene, and, in some cases, a gene predicted to encode alcohol dehydrogenase (**Figure 3**).

**Figure 3.**
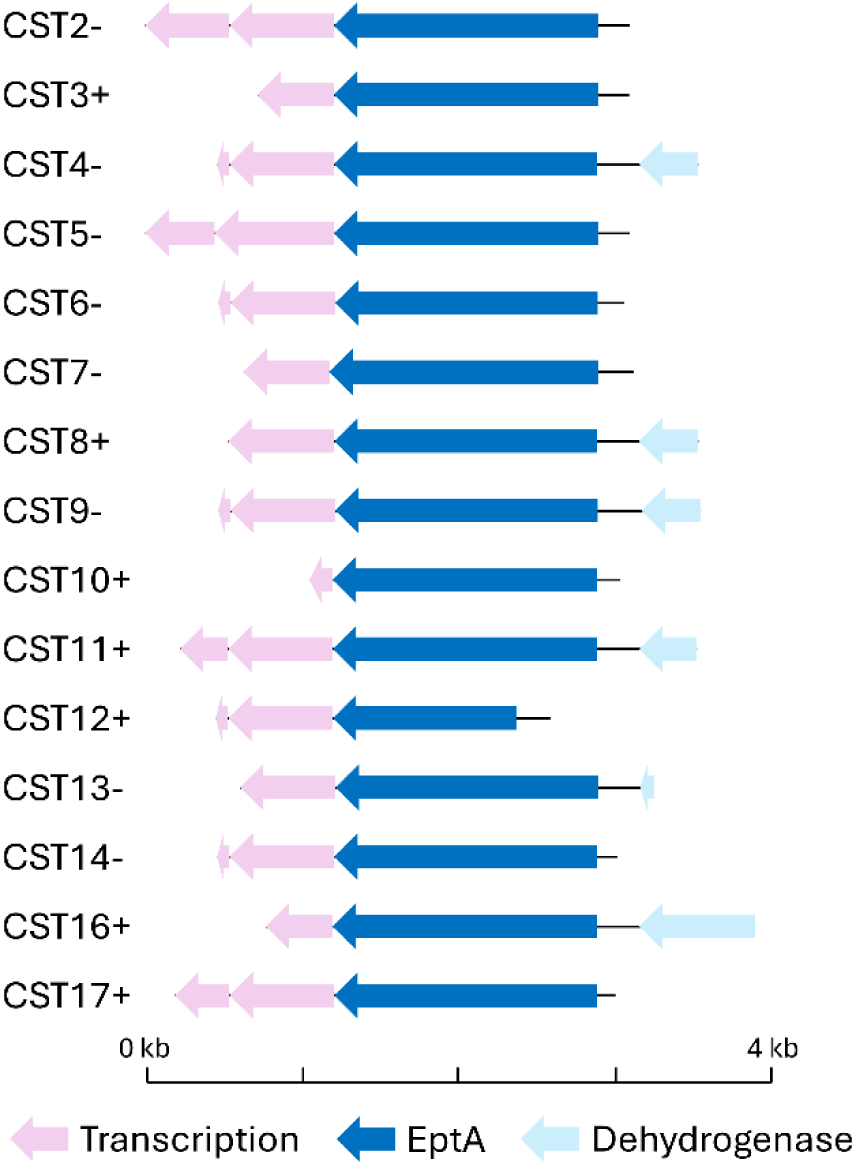
Gene schematics of colistin resistance-conferring metagenomic DNA fragments. Predicted open reading frames from DNA selected by 4 μg/ml (CST2-16) or 8 μg/ml (CST17) colistin selections. Annotations of predicted genes or their fragments include those likely to be related to transcriptional regulation (pink), *eptA* phosphoethanolamine transferase genes (blue), and a predicted alcohol dehydrogenase gene (light blue). The original orientation of the cloned *eptA* gene with respect to the vector promoter is denoted by ‘+’ or ‘-‘ next to the fragment name.

Using the predicted sequence of the EptA enzyme from the metagenomic fragment selected by growth on 8 μg/ml colistin (EptA17 from CST17), we prepared a phylogenetic tree containing homologs of EptA, MCR, EptB, and EptC (**Figure 4**). EptA17 clusters with the EptA protein from *Acinetobacter stercoris* and, more generally, with the branches of the tree mostly associated with EptA and MCR proteins.

**Figure 4.**
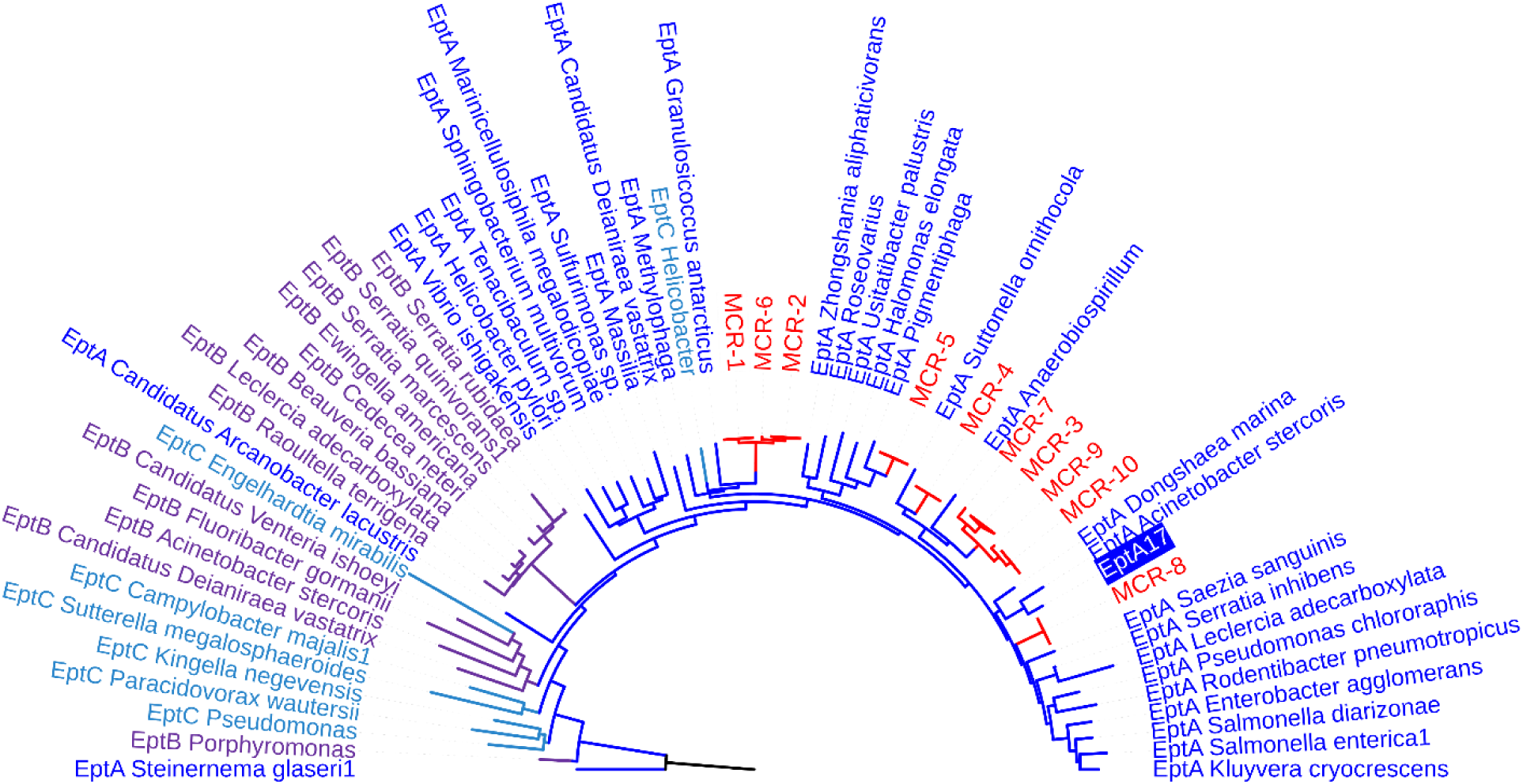
EptA, EptB, EptC, and MCR protein phylogenetic tree. Representative phosphoethanolamine transferase protein sequences from UniRef (EptA, EptB, EptC) or CARD (MCR) and the predicted EptA from metagenomic DNA fragment CST17 (EptA17). EptA sequences are shown in blue, EptB in purple, EptC in cyan, and MCR in red. EptA17 is highlighted with a blue background.

### *Acinetobacter* sp. *eptA* expression in *E. coli* confers clinically relevant levels of colistin resistance

We next performed agar dilution colistin susceptibility testing of the 15 functional metagenomic clones and a negative control *E. coli* strain. Except for CST4 and CST9, all of the clones with *eptA*-containing metagenomic gene fragments showed a decrease in colistin susceptibility compared to the plasmid-only control (**Figure 5**). Four clones, CST3, CST10, CST14, and CST17, grew at colistin concentrations greater than or equal to 2 μg/ml, with clone CST17 in particular showing robust growth at up to 4 μg/ml colistin. Of the clones showing the highest colistin resistance, three (CST3, CST10, and CST17) maintain their *eptA* gene in the same direction of the plasmid promoter and one (CST14) is oriented in the opposite direction (**Figure 3**).

**Figure 5.**
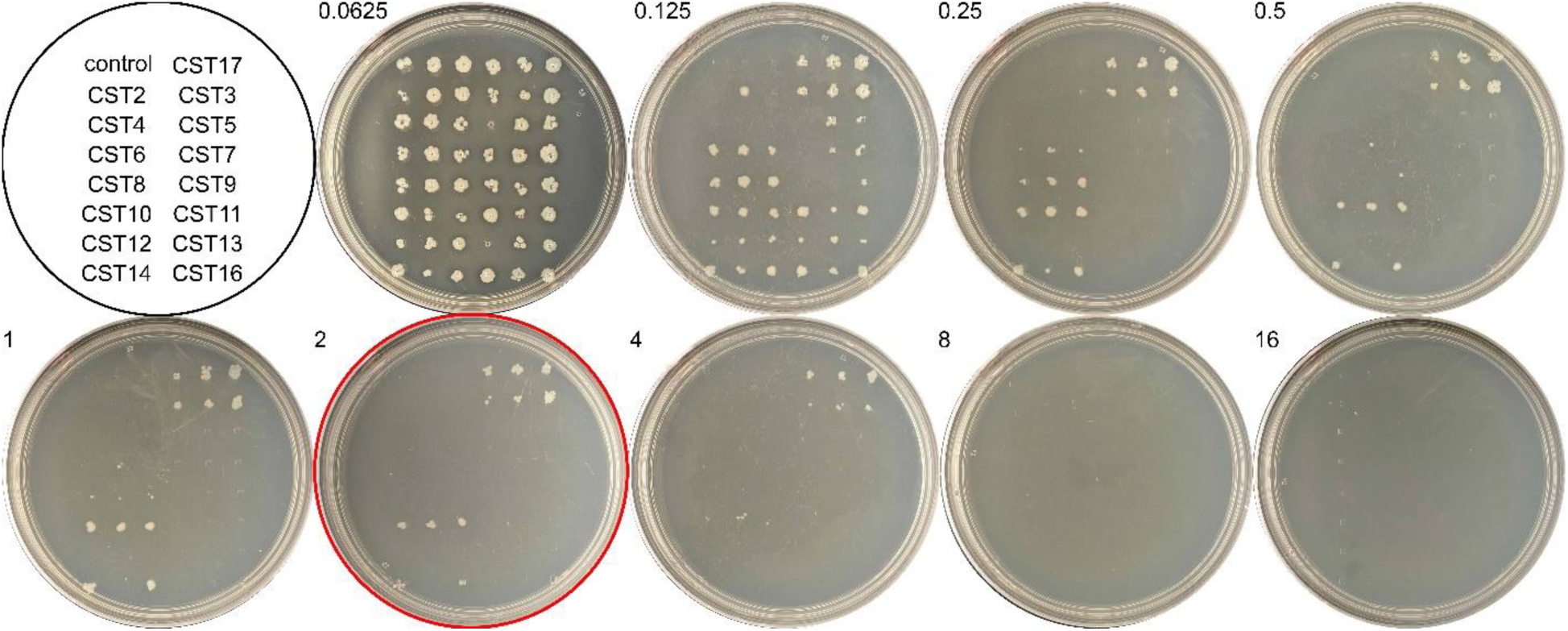
*eptA-*containing metagenomic DNA from *Acinetobacter* sp. decreases colistin susceptibility in *E. coli*. Agar dilution assay plates with 15 *eptA*-containing clones (CSTs) and one plasmid control (control) spotted in triplicate on agar plates containing colistin at the indicated concentrations (μg/ml). The 2 μg/ml plate is highlighted with a red circle.

We next decided to investigate colistin resistance of two of the *E. coli* clones from the selection. CST17 (with *eptA*17 encoded on its captured metagenomic DNA fragment), because it originated from a more stringent 8 μg/ml colistin selection, and CST10, because its relatively short metagenomic DNA fragment is almost completely composed of the predicted *eptA* gene (**Figure 3**). The two clones, alongside an empty plasmid negative control and an *mcr*-1-expressing positive control, were assayed by microbroth dilution (**Figure 6A**). The dose-response curves were solved to determine 50% inhibitory concentration (IC_50_) values (**Figure 6B**). The functionally selected clones were shown to have IC_50_ values of approximately 2.5 μg/ml and 4 μg/ml, similar to each other and the positive control. A one-way ANOVA test to compare these IC_50_ values against that of the negative control (∼1 μg/ml) showed that both functionally selected clones and the positive control are significantly more resistance to colistin than the negative control (p≤0.0005) (**Figure 6B**).

**Figure 6.**
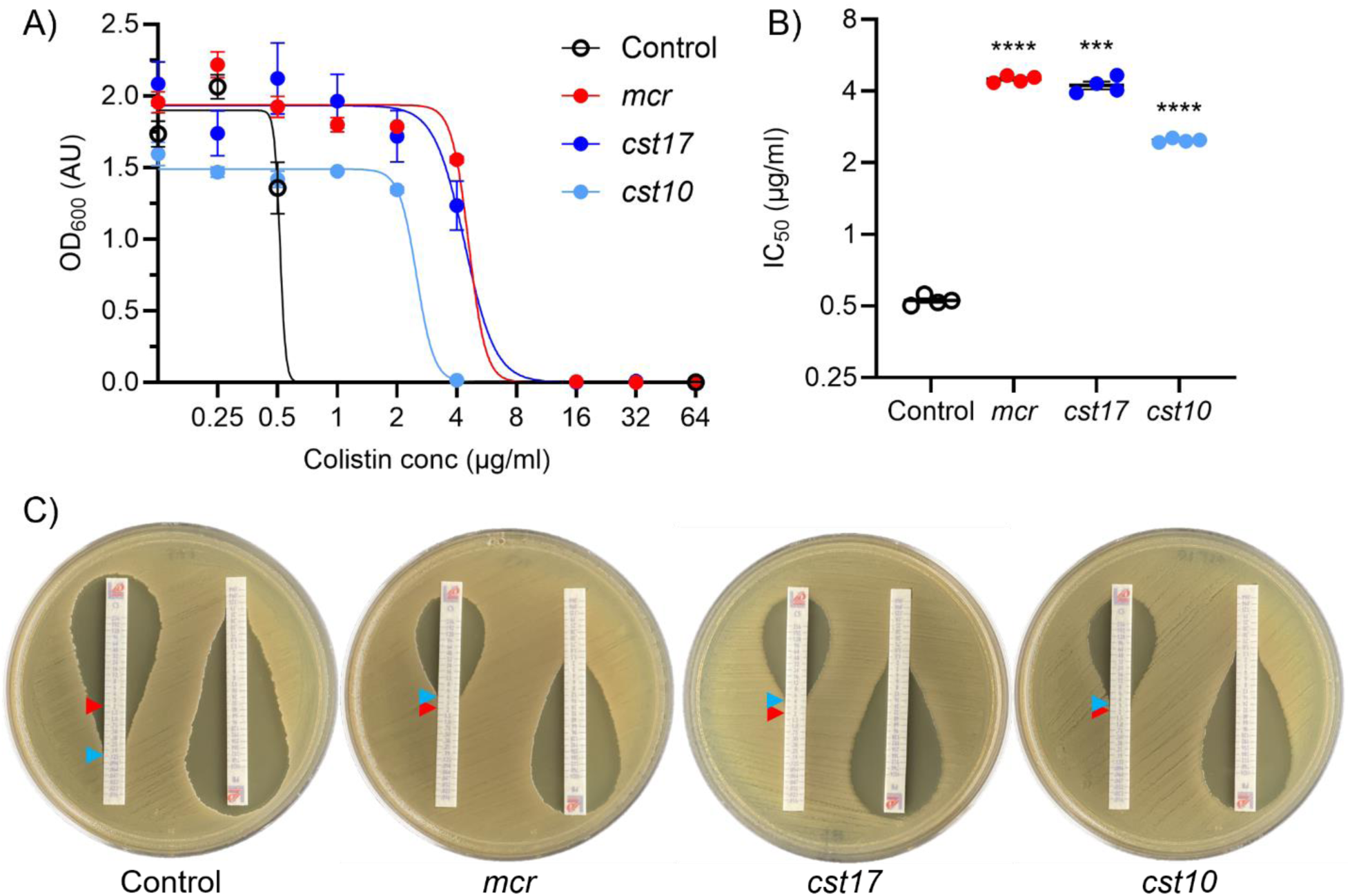
CST10 and CST17 confer colistin resistance in *E. coli*. A) Dose-response curve showing growth of *E. coli* harboring an empty plasmid (control, black) or plasmids containing functionally selected colistin resistant metagenomic DNA fragments (CST10 and CST17, light and dark blue) or a *bona fide mcr-1* gene (*mcr*, red). B) 50% inhibitory concentration (IC_50_) values calculated from the dose-response curves for the same strains (*** p = 0.0005 and **** p ≤ 0.0001 compared to control). For A) and B) n=4 replicates with standard error bars. C) Minimal inhibitory concentration (MIC) test strip assays for colistin (left strip) or polymyxin B (right strip) for the indicated strains. Red arrows indicate the 2 μg/ml breakpoint for colistin resistance and blue arrows indicate the observed MIC.

Finally, we determined minimal inhibitory concentration (MIC) values for colistin (and the related antibiotic polymyxin B) using MIC test strips on agar plates (**Figure 6C**). Clones CST10 and CST17 were able to grow in the presence of 3 μg/ml and 4 μg/ml colistin respectively, similar to the positive control at 4 μg/ml (**Figure 6C**). In contrast, growth of the negative control was inhibited by 0.25 μg/ml colistin (**Figure 6C**). Similar trends, but with lower MIC values, occurred with polymyxin B, showing resistance across polymyxin classes.

### Expression of functionally selected metagenomic DNA fragments leads to lipid A remodeling consistent with colistin resistance

EptA and MCR-mediated resistance to colistin in bacteria is due to modification of lipid A in the outer membrane (addition of a phosphoethanolamine moiety) that reduces interaction with the antibiotic (**Figure 1**). We next set out to verify that the observed colistin resistance in the CST10 and CST17 clones (**Figures 5 and 6**) was mediated by this mechanism. Clones CST10 and CST17, as well as the plasmid-only negative control and *mcr-1*-expressing positive control, were grown in liquid culture and harvested for their lipid A content. We used matrix-assisted laser desorption/ionization time-of-flight mass spectrometry to analyze mass changes in lipid A. The negative control, *E. coli* with an empty vector, produced a lipid species with mass consistent with the expected wildtype 3-deoxy-D-*manno*-octulosonic acid-lipid A (1797.2 m/z) (**Figures 7A and B**) (59). In contrast, the positive control *mcr-1*-expressing *E. coli* clone (**Figure 7B**), as well CST10 and CST17 (**Figures 7C and 7D**) contained an additional lipid with a mass increase of ∼123 m/z (1920.3 m/z), consistent with addition of a phosphoethanolamine group (**Figure 7A**) (59).

**Figure 7.**
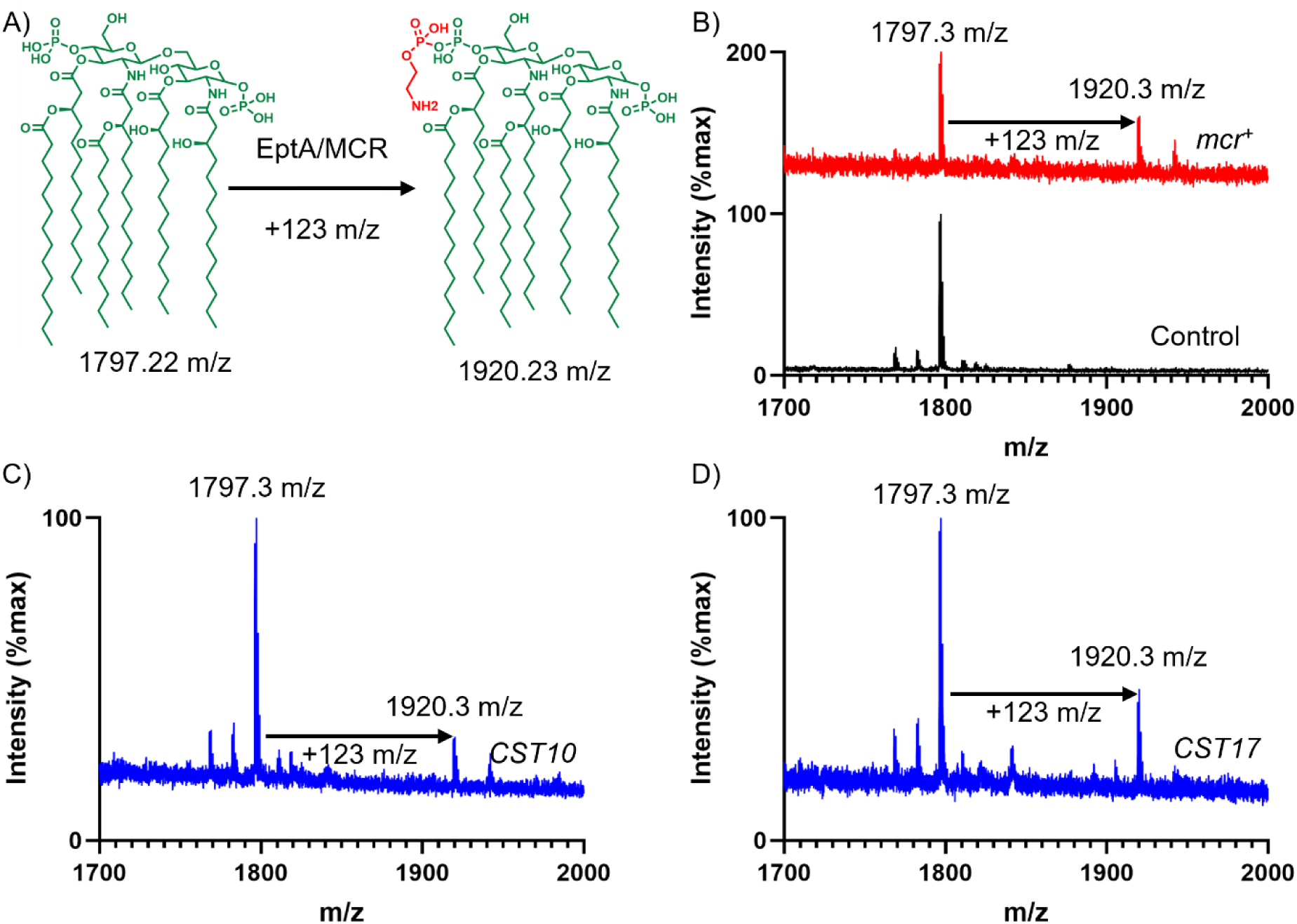
Expression of either *mcr* or CST genes leads to remodeling of lipid. **A.** A) Molecular reaction catalyzed by EptA and MCR: transfer of a phosphoethanolamine group (red) on to lipid A, resulting in a mass increase of 123 m/z. B) Mass spectrometry profile of lipids extracted from *E. coli con*taining either an empty plasmid vector (control, black) or a plasmid-expressed *mcr* gene (*mcr^+^*, red). Mass spectrometry profiles of *E. coli* expressing *eptA* genes from C) CST10 or D) CST17 agenomic DNA fragments.

### *eptA*17 shows evidence of mobilization within *Acinetobacter*

Because all the functionally selected colistin-resistant metagenomic DNA fragments appear to have the same origin (**Figure 3**), we used sequence information from CST16 and CST17 to construct a composite ‘full length’ sequence. As before (**Supplemental table 1**), this composite sequence has high nucleotide identity (>90%) to *Acinetobacter* species. We noticed, however, that the 5’ end and 3’ end of the composite sequence had highest identity to different *Acinetobacter* strains (this held true when we examined just CST17 by itself as well). Specifically, the 5’ end (including the *eptA* gene) aligns best to the chromosome of *Acinetobacter* species ASP199 (∼95% identity). This region also showed high sequence identity (∼94%) to the pAR3 plasmid from *A. radioresistens* strain DD78 (which contains a non-homologous toxin/antitoxin gene pair 3’ to the *eptA* region) (**Figure 8**). In both cases the 3’ region past the *eptA* gene of the composite sequence has essentially no homology locally to either *Acinetobacter* sp. ASP199 or the strain DD78 plasmid. Instead, the 3’ region (containing a fragment of a predicted alcohol dehydrogenase gene) has highest homology to another *Acinetobacter* strain, sp. XS-4. Here, sp. XS-4 only has local homology to the alcohol dehydrogenase gene (∼80% nucleotide identity) and does not encode an *eptA* gene in the vicinity (**Figure 8**).

**Figure 8.**
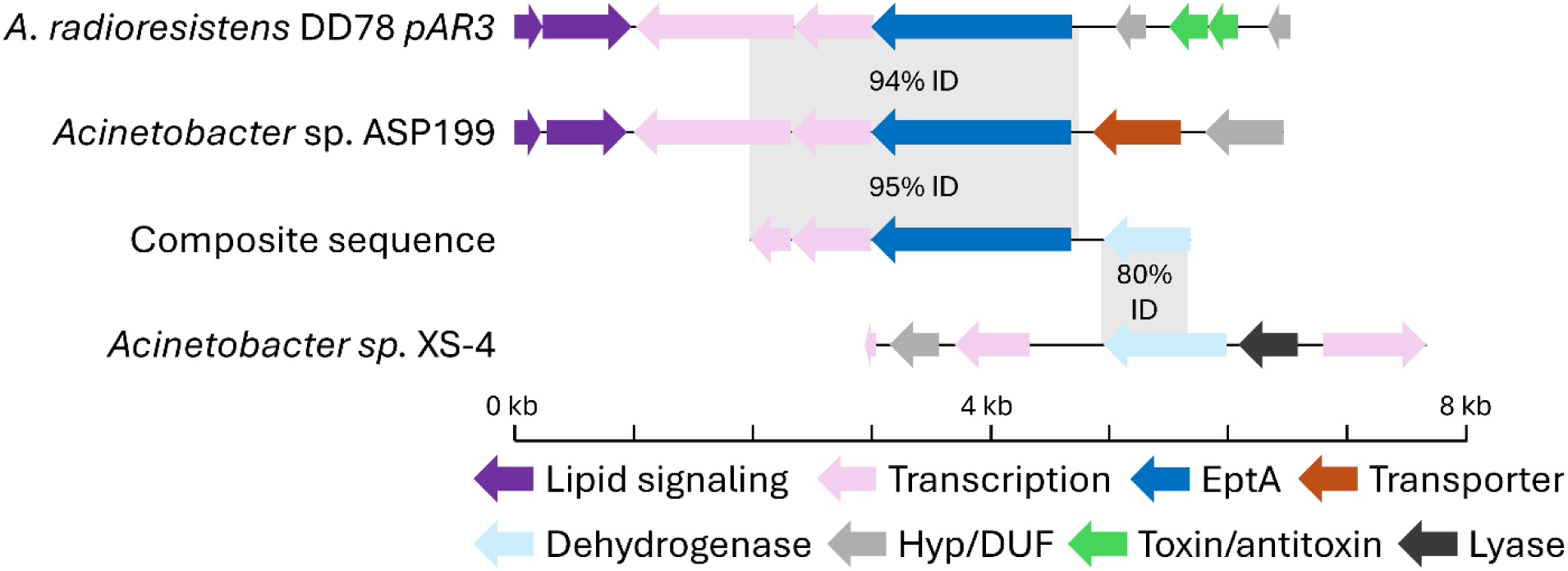
Potential genomic and plasmid native contexts for *eptA17*. Gene schematic representation for a composite sequence of the functionally selected metagenomic DNA fragments (‘composite sequence’, center). On top, gene schematics for genomic (*Acinetobacter* sp. ASP199) and plasmid (*A. radioresistens* DD78 pAR3) sequences with high nucleotide identity and coverage of the *eptA*-containing 5’ end of the composite sequence. On bottom, a gene schematic for genomic DNA from *Acinetobacter* sp. XS-4 with moderately high nucleotide identity and coverage of the 3’ end of the composite sequence containing an alcohol dehydrogenase gene fragment.

## DISCUSSION

We explored, in detail, colistin resistance determinants from a prior 4 μg/ml colistin functional metagenomic selection of a goose microbiome and a new 8 μg/ml colistin selection. Surprisingly, all 15 metagenomic gene fragments selected for study appear to originate from a single source, based on high nucleotide identity (**Supplemental figure 2**) and shared gene structure (**Figure 3**). Because the colistin-resistant gene fragments came from a small insert functional metagenomic library (the largest captured insert measures approximately 3 kb in length), we were unable to perfectly identify its original host genome. It is highly likely that the origin of this DNA fragment was an *Acinetobacter* bacterium (**Supplemental table 1**), but our analyses suggest that multiple *Acinetobacter* taxa may fulfill this role (**Figure 8**). The specific, local, context of the colistin-resistant insert is also unclear, with high nucleic acid identity regions mapping to both *Acinetobacter* chromosomal sequences and plasmid sequences (**Figure 8**). The homolog-containing plasmid (*A. radioresistens* DD78 plasmid pAR3, NCBI reference sequence NZ_CP038025.1), contains multiple predicted transposases and genes predicted to encode toxin-antitoxin proteins, suggesting mobilization of the plasmid as well as its cargo. The plasmid is also annotated as containing genes encoding metal resistance proteins (*e.g., terC*) and the CARD resistance gene identifier tool (51) identified a potential carbapenemase on the plasmid, highlighting that our captured phosphoethanolamine transferase gene may be part of a plasmid conferring resistance to antibiotics and other environmental pressures.

At the sequence, structural, and activity levels, EptA and MCR enzymes are almost indistinguishable (27, 30, 60). Both MCR and EptA enzymes are phosphoethanolamine transferases, and we confirmed the presence of this resistance mechanism in our own strains (**Figure 7C and 7D**). The essential distinction between *eptA* and *mcr* genes is their mobilization, or lack thereof, and their ability to confer colistin resistance (30). The European Society of Clinical Microbiology and Infectious Diseases Committee on Antimicrobial Susceptibility Testing (EUCAST) defines colistin resistance in *E. coli* as growth in the presence of ≥2 µg/ml colistin. We found that, in the cases of CST10 and CST17, carriage of the *eptA* gene studied in this manuscript confers colistin resistance past this level in *E. coli* (**Figure 6**). This fact, alongside very close relatives of this *eptA* gene having different genomic contexts across *Acinetobacter* taxa and appearing on a plasmid, suggests it may be appropriate to consider it an *mcr* gene, or at least a proto-*mcr* gene. While colistin resistance in the context of our study may be driven by the plasmid promoter upstream of the metagenomic fragment, it is notable that we found the *eptA*-containing DNA both in positive and negative reading frames with respect to this promoter (**Figure 3**). This suggests that in at least one of the cases where we observed growth in the presence of 2 μg/ml colistin (CST14, **Figure 5**), transcription was driven by the *eptA* gene’s native promoter and not the artificial promoter on our vector. Not all of the CST constructs supported more growth in the presence of colistin compared to the plasmid-only control strain. CST12 appears to have a truncation overlapping with the predicted *eptA* gene (**Figure 3**) and only supported growth in the presence of 0.125 μg/ml colistin compared to 0.0625 μg/ml for the control which is unlikely to be a significant difference. A few other CST constructs showed similarly negligible changes in colistin susceptibility. In *E. coli*, carriage of *mcr-1* comes with a metabolic burden, with intermediate levels of expression resulting in optimal antibiotic resistance (61). We hypothesize that the CST constructs that conferred no or limited changes in colistin susceptibility might be the result of mutations acquired during post-selection PCR amplification of resistance-conferring metagenomic DNA fragments (**Figure 1**, step 6). A future experiment cloning these highly similar, but apparently functionally different, *eptA* open reading frames could shed light on the potential balance between phosphoethanolamine transferase activity, metabolic burden, and colistin resistance.

Our functional metagenomic selection specifically captured a single *Acinetobacter eptA* gene rather than a collection of *eptA* homologs from other taxa, suggesting that *Acinetobacter* sp., a taxon well known for its high level of horizontal gene transfer and genomic plasticity (62), may be a source of *mcr* genes. This highlights one advantage of functional metagenomics over sequencing-only methods for identifying antibiotic resistance genes, particularly in this case where MCR enzymes appear the same as EptA enzymes phylogenetically (**Figure 4**). It has been shown that *Acinetobacter eptA* genes and their regulatory two-component signaling systems can be picked up by and/or induced by transposases, resulting in colistin resistance (63, 64, 64, 65). This observation parallels the discovery of *mcr*-1, as it was determined that it first emerged on a composite ISApl1 transposon that then transferred to plasmids of pathogenic bacteria (66). While *mcr*-1 was discovered on a swine farm (67), it is hypothesized that it was acquired from a natural environmental source, the antibiotic resistome (32). Migratory birds that can travel long distances are potential vectors of antibiotic resistance genes between antibiotic resistomes, and have the potential to disperse resistance genes along their flight path and destination (68–71). Antibiotic resistance genes found in gull feces have displayed evidence of horizontal transfer and diversification throughout the wildlife population, likely facilitated by the abundance of sewage, landfills, and public beaches encountered by gulls (69). Geese, too, have been implicated in being dispersal agents of antibiotic resistance genes through their migration paths (68, 72). In the specific case of *mcr* genes, in one study out of a sample of hundreds of birds approximately 50% carried the *mcr*-1 gene (70), while another study focused on the bar-headed goose in China found 7.3% of fecal samples contained colistin-resistant *E. coli*, often carrying *mcr-1* genes in the context of multi-drug resistant plasmids (71). These and our own results highlight the role bird microbiomes play as vectors for *mcr* or proto-*mcr* genes.

## Acknowledgments

CRediT contributions: Conceptualization – E.P.B and T.S.C.; Data curation – E.P.B. and T.S.C.; Formal analysis – E.P.B., Y.S., E.F., and T.S.C.; Funding acquisition – T.S.C.; Investigation – E.P.B., Y.S., E.F., and T.S.C.; Project administration – T.S.C.; Resources – T.S.C.; Supervision – T.S.C.; Visualization – E.P.B. and T.S.C.; Writing – original draft – E.P.B.; Writing – reviewing and editing – E.P.B., Y.S., and T.S.C. See Supplemental Figure 3 for graphical representation of contributions.

This work is supported by the Hatch Act of 1887, project award no. 7004080, from the U.S. Department of Agriculture’s National Institute of Food and Agriculture. The Bruker UltrafleXtreme MALDI TOFTOF mass spectrometer used by the School of Chemical Science Mass Spectrometry core was purchased in part with a grant from the National Center for Research Resources, National Institutes of Health (S10 RR027109 A).

## Data availability

Sequence data for CST2, CST3, CST4, CST5, CST6, CST7, CST8, CST9, CST10, CST11, CST12, CST13, CST14, CST16, and CST17 are available on GenBank.

